# Early Fibrin Biofilm Development in Cardiovascular Infections

**DOI:** 10.1101/2024.09.02.610803

**Authors:** Safae Oukrich, Jane Hong, Mariël Leon-Grooters, Wiggert van Cappellen, Johan A. Slotman, Gijsje H. Koenderink, Willem J.B. van Wamel, Moniek P.M. de Maat, Klazina Kooiman, Kirby R. Lattwein

**Author notes:** **Corresponding author:** Safae Oukrich, Erasmus MC, Biomedical Engineering, Dept. of Cardiology, Office Ee2322, P.O. Box 2040, 3000 CA, Rotterdam, the Netherlands. **Email addresses:**.

## Abstract

The single most common microbe causing cardiovascular infections is *Staphylococcus aureus* (*S. aureus*). *S. aureus* produces coagulase that converts fibrinogen to fibrin, which is incorporated into biofilms. This process aids in adherence to intravascular structures, defense against the host immune system, and resistance to antimicrobial treatment. Despite its significance, fibrin formation in *S. aureus* biofilms remains poorly understood. Therefore, this study aimed to elucidate the early development of cardiovascular biofilms. Clinically isolated coagulase-positive *S. aureus* and coagulase-negative *Streptococcus gordonii* (*S. gordonii*) from patients with cardiovascular infections, and a coagulase mutant *S. aureus* Δcoa, were grown in tryptic soy broth (TSB), Iscove’s Modified Dulbecco’s Medium (IMDM), and pooled human plasma, with or without porcine heart valves. Bacterial growth, metabolic activity, and bacterial fibrinogen utilization were measured over 24 hr at 37 °C. Time-lapse confocal microscopy was used to visualize and track biofilm development. *S. aureus* exhibited more growth in TSB and human plasma than *S. gordonii* and *S. aureus* Δcoa, but showed similar growth as *S. aureus* Δcoa in IMDM. Peak metabolic activity for all isolates was highest in TSB and lowest in human plasma. The presence of porcine valves caused strain-dependent alterations in time to peak metabolic activity. Confocal imaging revealed fibrin-based biofilm development exclusively in the coagulase-producing *S. aureus* strains. Between 2 and 6 hr of biofilm development, 74.9% (p=0.034) of the fibrinogen from the medium was converted to fibrin. Variations in fibrin network porosity and density were observed among different coagulase-producing *S. aureus* strains. Fibrin formation is mediated by *S. aureus* coagulase and first strands occurred within 3 hr for clinical strains after exposure to human plasma. This study stresses the importance of experimental design given the bacterial changes due to different media and substrates and provides insights into the early pathogenesis of *S. aureus* cardiovascular biofilms.

**Highlights:**

- Bacterial growth and activity are medium and substrate dependent
- Coagulase is necessary for *Staphylococcus aureus* fibrin biofilm development
- Fibrin strands begin forming in *Staphylococcus aureus* biofilms within 3 hours

## 1. Introduction

Cardiovascular bacterial infections, such as infective endocarditis (IE), which occur within the heart, and infections involving indwelling cardiovascular devices, like pacemakers and left ventricular assist devices, are life threatening. IE has a 1-year mortality rate of up to 50% [1], and cardiovascular device infections (CDI) have mortality rates ranging from 25-50%, depending on the device [2–5]. The most common infecting bacterium is *S. aureus*, responsible for at least 28% of IE and 54% of CDI [1, 6]. It is associated with prolonged hospital stays, higher mortality risk, and greater likelihood of surgical intervention needed to resolve the infection [7, 8]. Standard treatment for IE and CDI involves intensive, prolonged antibiotic therapy, which is often ineffective because these are biofilm infections [9]. Bacteria within biofilms encase themselves in an extracellular matrix, which makes them at least 10 to 1,000 times less susceptible to antibiotics than non-encased bacteria [10], and even 4,000 times for cardiovascular-related biofilms [11]. Despite the important role that biofilms have on increasing treatment complexity and mortality, the mechanisms underlying their early formation within the cardiovascular system are still not well understood.

Cardiovascular infections are thought to arise from bacteria interacting with pre-existing damaged or inflamed endothelial cells or by adhering directly to devices [12]. It is widely hypothesized that the host coagulation is activated upon bacteria entering the bloodstream [6, 13]. Staphylococci, streptococci, and enterococci cause at least 79% of cardiovascular infections [6]. These bacteria all express proteins that bind fibrinogen, however, only staphylococci produce enzymes to also convert fibrinogen to fibrin [14–16]. *S. aureus* can induce coagulation via two coagulases, staphylocoagulase and von Willebrand factor binding protein (vWbp), which both can be surface-bound and secreted by bacteria. These coagulases bind directly to prothrombin, bypassing the standard host coagulation cascade, thereby inducing a conformation change and activating prothrombin, triggering the conversion of fibrinogen into fibrin [6, 17]. Overall, little is known about the early formation and utilization of bacterial-induced fibrin in cardiovascular biofilms.

Human plasma is a more physiologically accurate growth medium to replicate cardiovascular biofilms compared to traditional bacterial or mammalian cell culture media, yet plasma alone is rarely utilized as a growth media. Commonly, studies have used varying concentrations of plasma supplemented with bacterial or mammalian cell culture media, such as up to 50% plasma in Brain-Heart Infusion (BHI) [18], Mueller Hinton, and Tryptic Soy broth [19, 20] 25% or 30% of plasma in 20% or 70% Roswell Park Memorial Institute medium (RPMI) [21], respectively. These studies demonstrated that the addition of plasma to the media enhances biofilm formation in a dose-dependent and isolate-dependent manner. For the few studies that used 100% human plasma, calcium chloride was added with the planktonic bacterial inoculation to artificially rapidly induce clot polymerization [22, 23]. In one study, fibrin biofilms have been grown on human-whole blood retracted clots in 100% plasma without calcium chloride, with only 24 hr end-point biofilm visualization being performed [24].

Here we aimed to understand the early development of fibrin biofilms in cardiovascular infections during the first 24 hr of its formation. To first understand the impact of media choice and the substrate bacteria grow on, such as whether the infection occurs on heart tissue versus a hard surface (e.g. an indwelling device), we compared the bacterial growth and metabolic activity of clinical IE isolates of *S. aureus* and *Streptococcus gordonii* (which binds fibrinogen but cannot convert it into fibrin), along with a coagulase mutant *S. aureus* (*S. aureus* Δcoa), using three different growth media: Tryptic Soy broth (TSB), Iscove’s Modified Dulbecco’s Medium (IMDM), and 100% human plasma. Fibrinogen utilization was measured during biofilm development and high-resolution fluorescence confocal microscopy was used to visualize biofilms grown in human plasma on porcine valves. Furthermore, time-lapse microscopy was used to track and analyze *S. aureus* fibrin formation in biofilms over time.

## 2. Materials and Methods

### 2.1 Bacterial isolates

*Staphylococcus aureus* (*S. aureus*) 25268, *S. aureus* 50825, and *Streptococcus gordonii* (*S. gordonii*) strains were isolated from patients diagnosed with IE at the Erasmus MC, the Netherlands and stored at −80 °C. According to institutional policy, bacterial isolates de-identified and anonymized are classified as non-human subject research and do not require informed consent. The coagulase mutant *S. aureus* strain (*S. aureus* Δcoa) was constructed from the widely-used wildtype lab strain 8325-4. *S. gordonii*, a coagulase-negative streptococcal strain, served as another control for *S. aureus* Δcoa. Before conducting experiments, *S. aureus* 25268*, S. aureus* 50825*, S. aureus* 8325-4 and *S. gordonii* were grown at 37 °C on 5% sheep blood agar plates (tryptic soy agar; BD, Trypticase^TM^, Thermo Fisher Scientific, Waltham, MA, USA). *S. aureus* Δcoa was grown on tryptic soy agar plates containing 5µg/ml tetracycline. *S. aureus* strains were incubated overnight, while *S. gordonii*, being since it is a slow-growing strain, was incubated for two nights.

### 2.2 Biofilm formation

Biofilms were formed *in vitro* by suspending single bacterial colonies from overnight plate cultures in 4 ml of 0.9% NaCl saline solution until an optical density (OD) of 0.5 (±0.05) was reached at 600 nm as determined using a cell density meter (Ultraspec 10, Amersham Biosciences, Little Chalfont, UK). The inoculated NaCl solution was added to growth medium to reach 10^6^ colony forming units per ml. Biofilms were grown in either Tryptic Soy Broth (TSB), Iscove’s Modified Dulbecco’s Medium (IMDM; 11500556; Gibco, Bleiswijk, the Netherlands), or 100% human plasma (blood type O^+^; pooled from 5 healthy donors and directly stored at −80 °C; anticoagulant citrate phosphate double dextrose; Sanquin, Rotterdam, the Netherlands). Before experiments, the growth medium was pre-warmed to 37 °C. Biofilms were cultured at 37 °C, with or without a 6 mm porcine valve punch biopsy in either sterile flat-bottom 96-well polystyrene tissue culture plates (CellStar; Greiner Bio-One, Alphen aan de Rijn, The Netherlands), CalWel inserts (SymCel, Solna, Sweden), 1.5 ml Eppendorf tubes, or tissue culture-treated µ-Slide mono-channels (80916; ibiTreat µ-Slide I0.8 Luer; Ibidi GmbH, Martinsried, Germany). Porcine hearts were acquired from a local butcher. The valve biopsies were excised, biopsied, and then pre-sterilized for 5 min in 0.25% glutaraldehyde (11433488; glutaraldehyde, 25% aqueous solution, Thermo Scientific Chemicals, Dublin, Ireland) and rinsed thoroughly. To investigate the influence of the substrate on biofilm development, porcine valve tissue was used to mimic native heart valve infections, while empty sample containers (wells, µ-Slides or inserts) were used to simulate non-native tissue infections. At least n=3 was performed for all experiments.

### 2.3 Biofilm growth and microcalorimetry

Growth and metabolic activity over time was assessed for *S. aureus* 25268, *S. aureus* Δcoa, and *S. gordonii* using TSB, IMDM, and human plasma. For growth measurements, 200 µl of inoculated medium was added to each well of a the 96-well tissue culture plate. The plate was then placed in a microplate reader (FLUOstar Omega; BMG LABTECH GmbH, Orternberg, Germany) to measure growth (OD_600_) every 15 min for 24 hr at 37 °C. Microcalorimetry was used to assess the heat produced in microWatts (µW) over time to determine bacterial metabolic activity using the CalScreener (SymCel, Solna, Sweden), a closed and isothermal system. This method enables the measurement of metabolic activity in the presence of porcine valves, which is not feasible with optical density measurements. Inoculated media (200 µl) was added to CalWell inserts which were then placed in titanium cups (SymCel, Solna, Sweden) and loaded into the CalScreener for 24 hr at 37 °C to determine heat flow over time.

### 2.4 Determination of bacterial fibrinogen utilization

Bacterial fibrinogen utilization was determined at 0, 1, 2, 4, 6, and 24 hr using the functional Clauss method [25] on the Sysmex CS-5100 system (Siemens, Den Haag, the Netherlands). The decrease in fibrinogen levels in the plasma over time reflects the binding of fibrinogen to the bacteria themselves or conversion into fibrin used to form the biofilms. Biofilms of *S. aureus* 25268, *S. aureus* 50825, *S. aureus* Δcoa, *S. aureus* 8325-4, and *S. gordonii* were grown in human plasma within Eppendorf tubes as described in section 2.2. Individual samples were prepared at each time point (n=3) and centrifuged at 14,000 g for 10 min. The supernatant was then transferred to a new Eppendorf and centrifuged again at 15,000 g for five min and stored at −80 °C until measuring fibrinogen levels.

### 2.5 Bacteria and fibrin visualization

All images were acquired using a custom-built, upright Nikon A1R+ confocal microscope [26]. Biofilms of *S. aureus* 25268, *S. aureus* Δcoa, *S. aureus* 8325-4 and *S. gordonii*, which were grown in human plasma on sterile porcine valve biopsies for 24 hr at 37 °C, were placed in a glass-bottom dish (D35-10-0-N, Cellvis; IBL Baustoff + Labor GmbH, Gerasdorf, Austria) for imaging. These biofilms were imaged using a 60× water-dipping objective (CFI Plan 60XC W, 2.5 mm working distance, NA 1.0, Nikon Instruments, Amsterdam, the Netherlands). For each porcine valve sample, three z-stacks were captured with a 2 µm step size, producing images of 212 µm x 212 µm at a resolution of 512 x 512 pixels. Time-lapse imaging was performed on *S. aureus* 25268, *S. aureus* 50825, and *S. aureus* 8325-4 grown in human plasma for the first 6 hr after inoculation (t=0) and again at 24 hr in Ibidi µ-Slide mono-channels. For these experiments, the confocal microscope was equipped with a 100× water dipping objective (CFI Plan 100XC W, 2.8 mm working distance, NA 1.1, Nikon Instruments, Amsterdam, the Netherlands). For each time-lapse image, a z-stack of 800 µm was captured at the same location of the slide (from top to bottom) was made with a 2 µm step size at 1-1.9 frames/sec, with each image at 128 µm x 128 µm (with 512 x 512 pixels). To visualize fibrin(ogen) within biofilms, human fibrinogen fluorescently-labelled with Alexa Fluor 647 (final concentration 0.1 mg/ml in plasma containing ∼ 2.0 mg/ml; F35200; Thermo Fisher Scientific; excited at 640 nm and detected at 700/75nm) was added to the plasma just before bacterial inoculation. Before imaging, SYTO^TM^ 9 green fluorescent nucleic acid stain (2 μl/ml; S34854; Thermo Fisher Scientific; excited at 488 nm and detected at 525/50 nm) was used to visualize living bacteria, and propidium iodide (PI; 25 μl/ml; P4864-10ML; Sigma-Aldrich, Zwijndrecht, the Netherland; excited at 561 nm and detected at 595/50 nm s), to visualize porcine valvular cells and dead bacteria. Amino sugars (N-Acetylglucosamine) in the biofilms grown on porcine valves were visualized using wheat germ agglutinin (WGA) labelled with Alexa Fluor^TM^ 350 (6 μl/ml W11263; Thermo Fisher Scientific; excited at 405 nm and detected at 425/75 nm).

### 2.6 Data analysis

Confocal images were obtained and analyzed using Nikon Instruments Software (NIS)-Elements Advanced Research (Version 4.50; Nikon Instruments). Additional post-hoc image analysis on confocal microscopy images was performed using a custom-designed ImageJ/Fiji [27] script (Optical Imaging Center, Erasmus MC, Rotterdam). Briefly, fibrin density was measured by segmenting the fibrin network by thresholding and determining the ratio of density of fibrin over the total imaged volume. The porosity was measured by segmenting the empty spaces inside the fibrin network by thresholding. Subsequently, the largest possible circle centered around the segmented spaces not containing any pixels above an intensity of 1700 was iteratively determined as the threshold. The porosity was calculated by taking the diameter of the fitted circles and calculating the surface. Graphical representation and data analysis (including baseline correction) was performed with Microsoft Excel (Microsoft Corporation, 2010), MATLAB (MATLAB, version 2024a), and SPSS (IBM SPSS Statistics, Version 28). Normality of the data was tested using the Shapiro-Wilk test. Statistical significance between groups was determined using Mann-Whitney-U test, and a paired t-test or Wilcoxen signed rank test for changes between time points within a group. A p-value < 0.05 was considered statistically significant. All quantifiable data points are presented as mean ± standard deviation (SD).

## 3. Results

### 3.1 Bacterial growth and metabolic activity is medium and substrate dependent

The effect of different growth media on bacterial growth rates were investigated for *S. aureus* 25268, *S. aureus* Δcoa, and *S. gordonii* over a 24 hr period. The bacteria were grown in TSB (Fig. 1A; top graph), IMDM (Fig. 1B; middle graph), and human plasma (Fig.1C; bottom graph). In TSB, *S. aureus* 25268 (blue line) demonstrated the highest growth rate, with *S. aureus* Δcoa (yellow line) lower and *S. gordonii* (green line) even lower growth rates in this medium. In IMDM, *S. aureus* Δcoa and *S. gordonii* were comparable to growth in TSB, with a slight delay in the growth of *S. aureus* Δcoa compared to *S. aureus* 25268. Despite this delay, the final OD values of *S. aureus* Δcoa and *S. aureus* 25268 after 24 hr were similar. In human plasma, *S. aureus* 25268 had the highest growth rate and final OD, indicating that human plasma was the most favorable medium for its growth. The growth of *S. aureus* Δcoa and *S. gordonii* remained lower than that of *S. aureus* 25268 across all three media, except in IMDM after 14 hr when both strains reached similar biofilm OD measurements.

**Figure 1.**
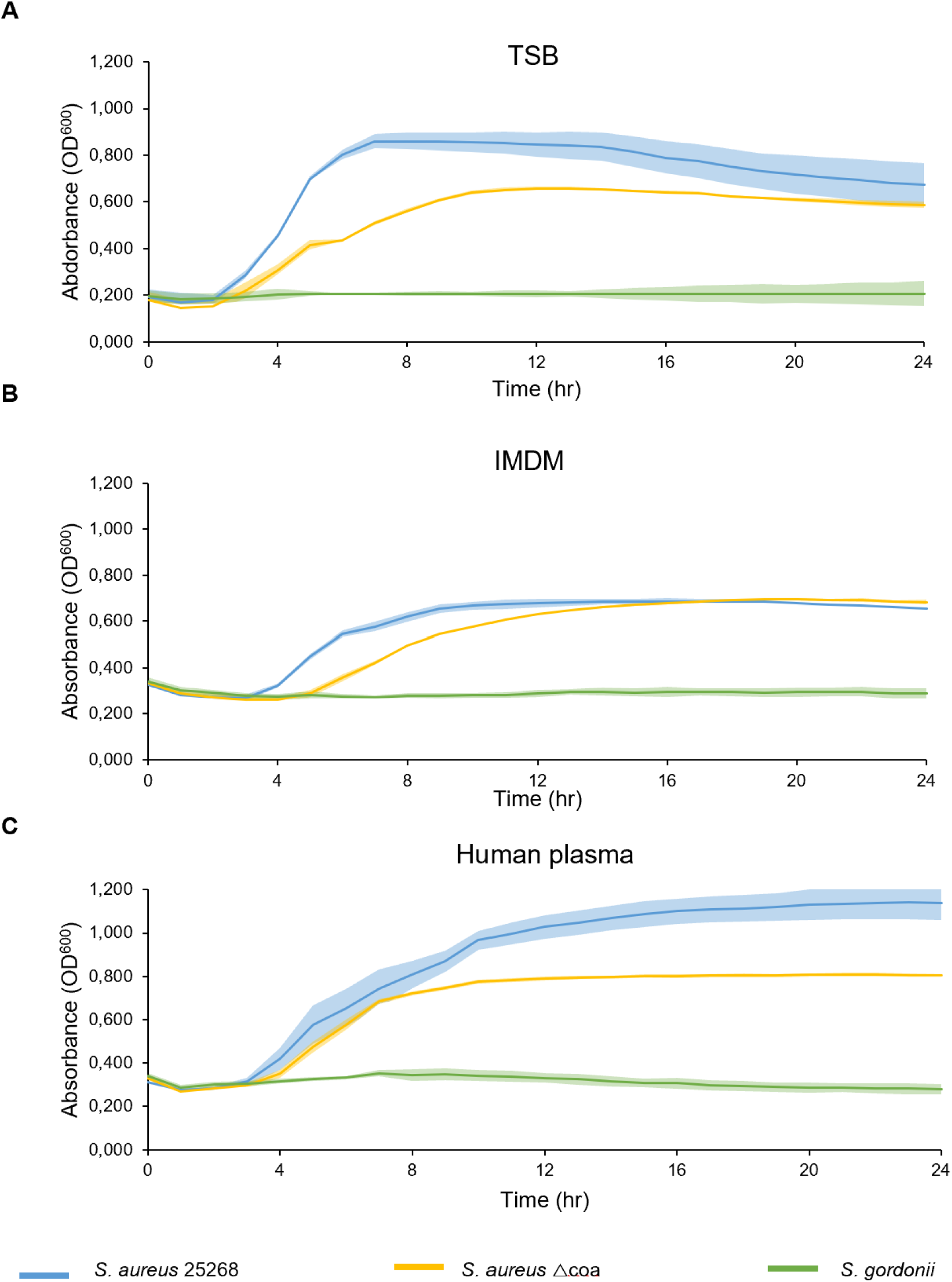
Bacterial growth curves of three different strains cultured in three different types of growth media. *S. aureus* 25268 (blue), *S. aureus* △coa (yellow), and *S. gordonii* (green) were grown in **(A)** tryptic soy broth (TSB), **(B)** Iscove’s Modified Dulbecco’s Medium (IMDM) and **(C)** human plasma. Bold singular lines represent the mean and the corresponding shadowing represents the standard deviation (n=4).

The metabolic activity, which includes growth as well as various other processes like virulence factor production (e.g., coagulase), of *S. aureus* 25268, *S. aureus* Δcoa and *S. gordonii* was assessed over 24 hr in the three different media: TSB (Fig. 2A), IMDM (Fig. 2B) and human plasma (Fig. 2C). Control experiments with sterile TSB, IMDM, and human plasma, both with and without porcine valve biopsies, exhibited no metabolic activity (data not shown). In TSB (Fig. 2A), *S. aureus* 25258 exhibited the highest metabolic activity, peaking at 60.99±1.6 µW (solid blue line). *S. aureus* Δcoa showed a similar peak, albeit lower at 43.8±0.7 µW (solid orange line). *S. gordonii* had even lower metabolic activity, peaking at 29.11±3.5 µW (solid green line). The addition of porcine valve material affected the metabolic activity of all strains differently (Fig. 2A, dashed lines). *S. aureus* 25268 showed reduced activity after 8 hr, *S. aureus* Δcoa exhibited increased activity from the time to peak until the 14 hr time point before declining, and *S. gordonii* had reduced overall activity but achieved its peak sooner. The time to reach peak metabolic activity in TSB was approximately 4.5-5 hr for the *S. aureus* strains, and 8-12.7 hr for *S. gordonii* (Fig. 3).

**Figure 2.**
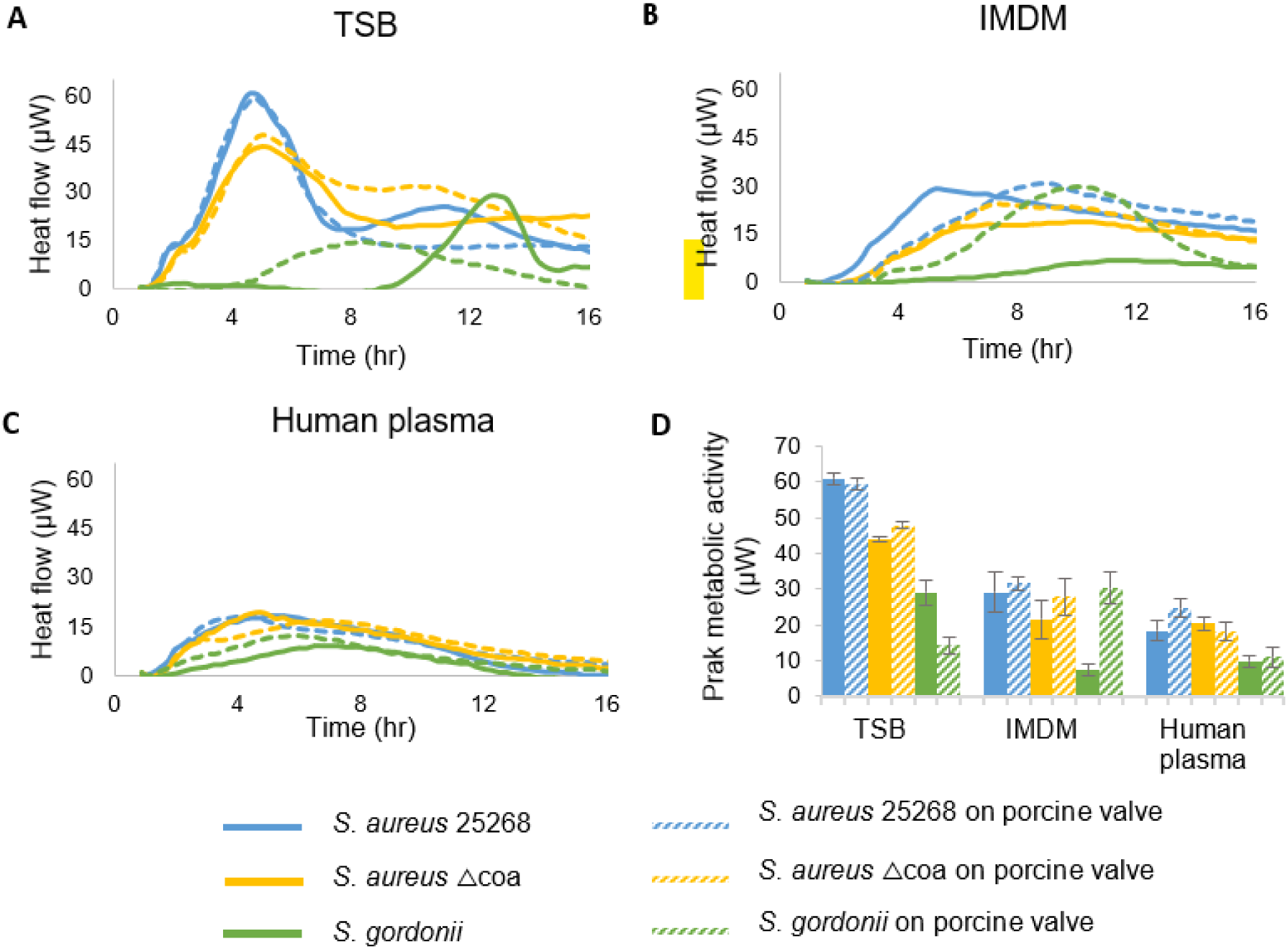
Total metabolic activity (µW) of **(A)** *S*. *aureus* 25268, **(B)** *S. aureus* △coa, and **(C)** *S. gordonii* measured in tryptic soy broth (TSB), Iscove’s Modified Dulbecco’s Medium (IMDM), and human plasma with (dashes lines) or without (solid lines) porcine valve biopsies**. (D)** Maximum metabolic activity of each strain. Lines and bars represent the mean. Error bars represent the standard deviation (n=3).

**Figure 3.**
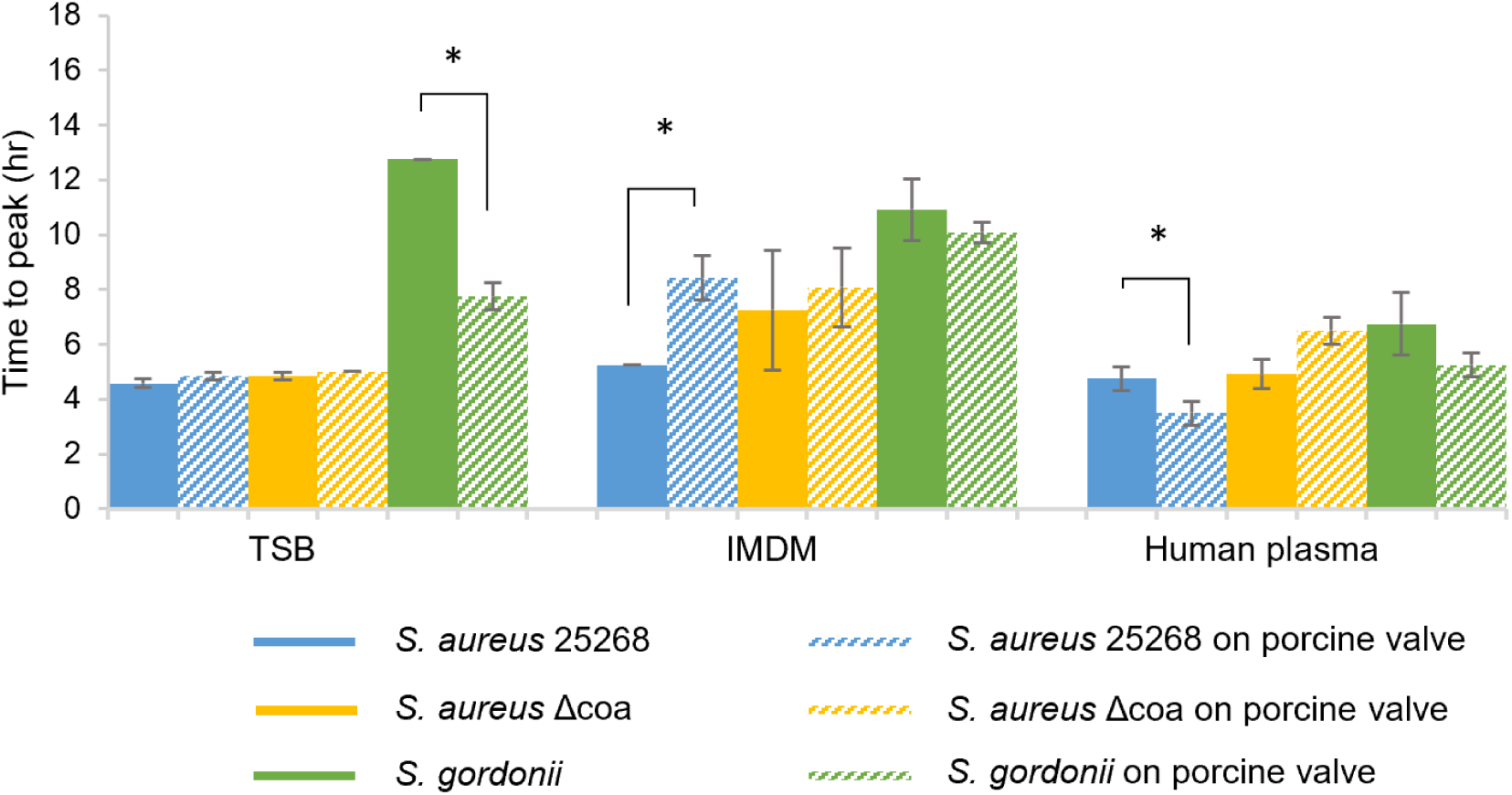
Time to peak metabolic activity of *S. aureus* 25268, *S. aureus* △coa, and *S. gordonii* measured in tryptic soy broth (TSB), Iscove’s Modified Dulbecco’s Medium (IMDM) and human plasma, with or without porcine valve present. A single asterisk denotes statistical significance (p < 0.05) between same strains with and without porcine valves. Data is presented as the mean and error bars the standard deviation (n=3).

In IMDM (Fig.2B), notable differences were observed compared to TSB for all strains. Without valve tissue, *S. aureus* 25258 peaked around 29.22±5.6 µW, *S. aureus* Δcoa at 21.5±5.4 µW, and *S. gordonii* at 7.22±1.6 µW. With porcine valve material, *S. aureus* 25258 took approximately 4 more hours to reach its peak; *S. aureus* Δcoa showed increased activity, peaking higher and lasting again until the 14 hr time-point; *S. gordonii* showed a distinct increase, with a 20.33±4.52 µW higher peak activity. It generally took longer to reach peak heat flow in IMDM compared to TSB for all strains, except *S. gordonii* without porcine valves which peaked 2.88±0.4 hours sooner (Fig. 3).

In human plasma (Fig. 2C), all strains exhibited the lowest metabolic activity of all three media, with similar heat flow pattern but different maximum heat flows achieved. The addition of porcine valve material again affected heat flow for all strains. For *S. aureus* 25258 in human plasma, it took less time to reach maximum activity without the porcine valve, but for *S. aureus* Δcoa maximum activity was achieved with the valve tissue present (Fig. 3). Overall, bacterial metabolic activity was found to be media- and substrate-dependent. *S. aureus* 25258 demonstrated the highest metabolic activity across all media and the presence of a porcine valve affected the heat flow and time to maximum metabolic activity, and significantly for *S. aureus* 25258 in IMDM and human plasma, and for *S. gordonii* in TSB (Fig. 3).

### 3.2 Only the coagulase-producing S. aureus strain developed fibrin biofilms

Confocal microscopy was used to visualize fibrin(ogen) presence within biofilms of different strains grown in plasma on porcine valves for 24 hr at 37°C (Fig. 4). Nuclei of porcine valvular cells and dead bacteria, both stained red, could be distinguished based on size difference, as valvular nuclei are oval at ± 10 µm wide and ± 4 µm long and dead bacteria are spherical at ± 1 µm in diameter.

**Figure 4.**
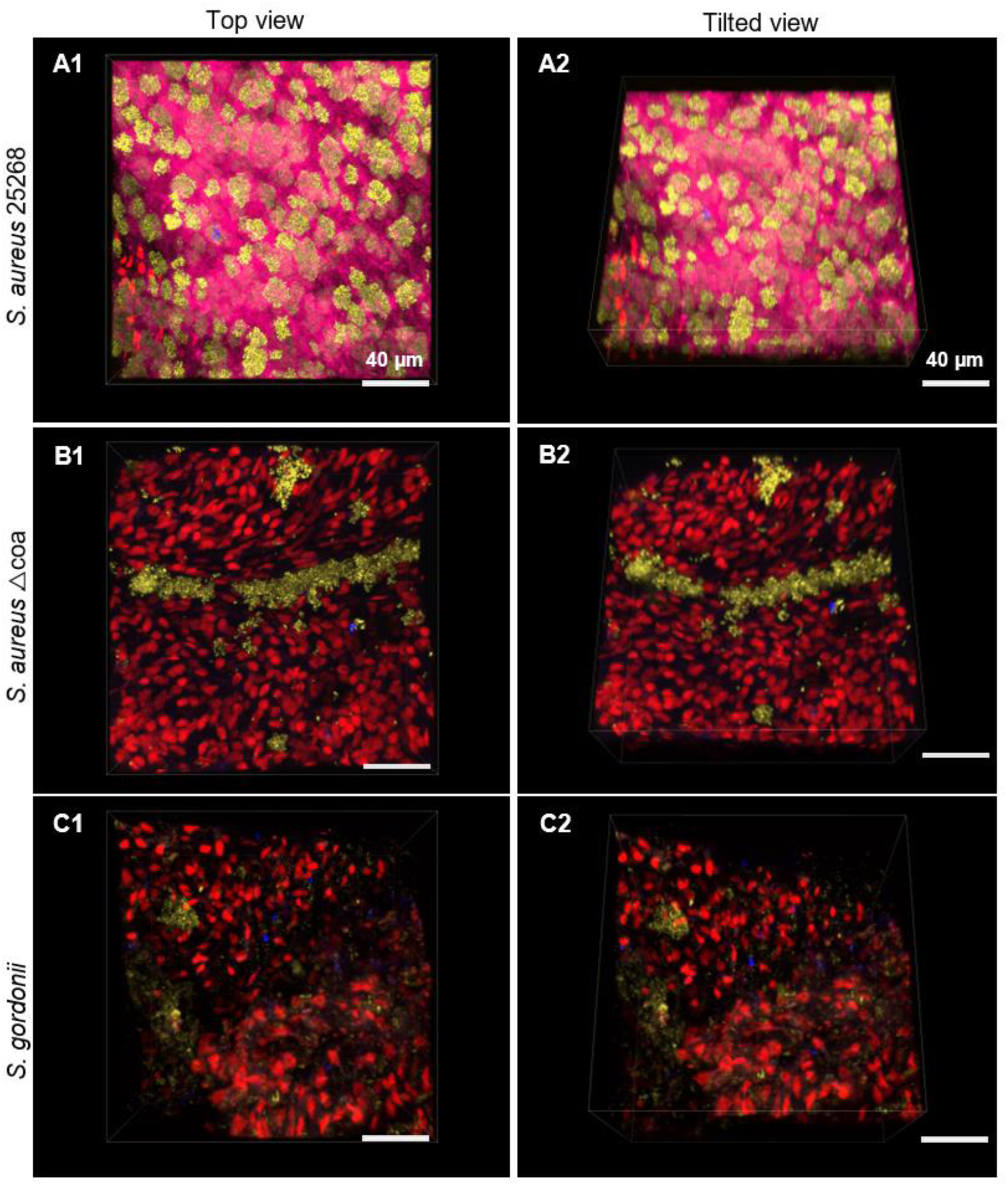
Confocal three-dimensional volume-rendered images of *S. aureus* 25268 25268 (**A**), *S. aureus* △coa (**B**), and *S. gordonii* (**C**) biofilms grown for 24 hr at 37 °C in human plasma on porcine valves. **A1**, **B1**, **C1** are the top views of biofilms, while **A2, B2, C3** are corresponding views at an angle. For all images, blue (Alexa Fluor 350 wheat germ agglutinin) indicates amino sugars, yellow (pseudo-colored; Syto9) indicates living bacteria, red (propidium iodide) indicates valvular cells and dead bacteria, and pink e (Alexa Fluor 647) indicates fibrin and fibrinogen. All images are representative of three separate experiments.

Plasma-grown *S. aureus* 25268 developed biofilms with a dense fibrin strand network (pink) covering the surface of the valve (Fig. 4 A1, 2). The majority of the bacteria were living (pseudo colored yellow) and formed aggregates approximately 15-35 µm in size within the fibrin strand network, with fibrin(ogen) found within the aggregates as well as covering the surface. In contrast, *S. aureus* Δcoa (Fig. 4 B1, 2) developed biofilms without any fibrin or fibrinogen and were dispersed over the porcine valve, while it’s wild-type strain, *S. aureus* 8325-4 (Fig. S1 A), was able to develop a fibrin network, albeit much less than the clinical *S. aureus* isolates. *S. gordonii* (Fig. 4 C1, 2) developed biofilms without a fibrin network and no fibrin strands were observed. Sparse fibrin(ogen) clumps were observed throughout the biofilm. The amount of amino sugars present appeared slightly more for *S. gordonii* than for the *S. aureus* strains.

### 3.3 Coagulases are necessary for S. aureus fibrinogen utilization

To further investigate the role of coagulases in the conversion of fibrinogen into fibrin, a fibrinogen utilization assay was performed. Figure 5 shows the percentage of fibrinogen used by bacteria, i.e., the decrease in fibrinogen measured, during 24 hr biofilm development. The starting concentration of fibrinogen in plasma was 2.0 g/l. *S. aureus* 25268 (blue line) showed a rapid increase in fibrinogen utilization, reaching 76.4 % within 6 hr, 78.1% at 8 hr, and then maintaining this level for the duration of the 24 hr period. *S. aureus* Δcoa (yellow line) exhibited minimal fibrinogen utilization, with only 5.1% maximally measured over 24 hr, which was consistent with the plasma-only control samples (red line). While *S. aureus* 8325-4 (purple line) reached 9.0% at 24 hr (Fig. S1 B). *S. gordonii* (green line) moderately bound fibrinogen, gradually reaching 21.3% at 9 hr and remained at this level for the remainder of the 24 hr. A significant drop in fibrinogen levels was observed for *S. aureus* 25268 between 2 and 6 hr, 74.9% (p=.034), indicating a sharp increase in fibrinogen utilization during this period. This was not observed in the coagulase-negative strains, *S. aureus* Δcoa and *S. gordonii* (10.2%) or in the plasma-only control samples.

**Figure 5.**
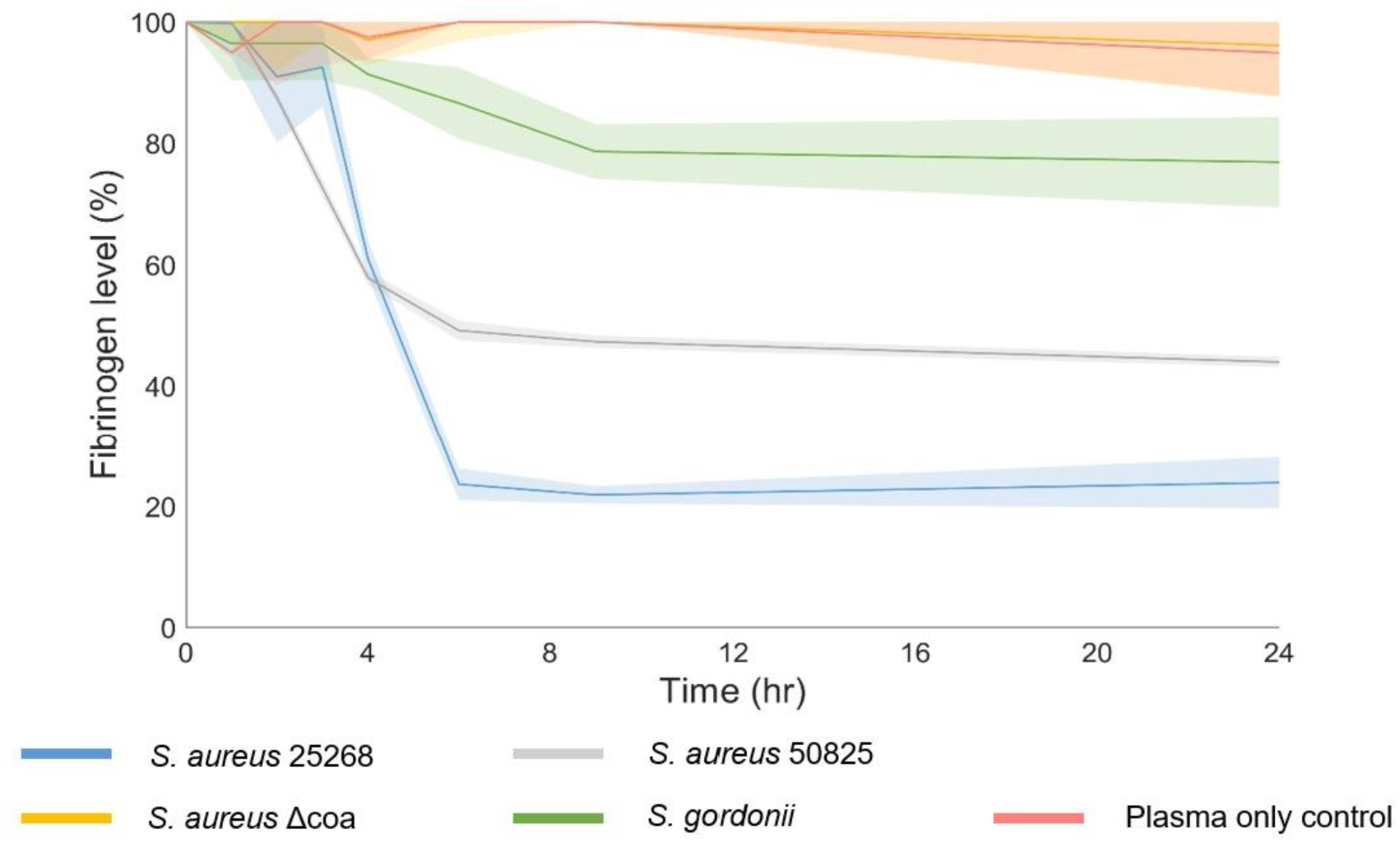
Bacterial fibrinogen utilization of *S. aureus* 25268 (blue), *S. aureus* △coa (yellow) and *S. gordonii* (green) over 24 hr. Human plasma alone (red) was used as a control. Solid lines represent the mean and shading the standard deviation of three independent experiments.

### 3.4 Fibrin network development occurs within the first 6 hours

The development of the fibrin network in biofilms formed by different strains of *S. aureus* were analyzed using time-lapse imaging. Figure 6 provides representative images of fibrin network development for *S. aureus* 25268 (6A) and *S. aureus* 50825 (6B). In each image, fibrin(ogen) (pink) was fluorescently-labeled to visualize the biofilm structure. Early time points (1-3 hr) revealed sparse and scattered bacterial cells covered in fibrin(ogen), while later time points (4-6 hr) quickly showed a progressive increase in density of fibrin strands and growing aggregates of bacteria within the biofilm matrix. *S. aureus* 25268 was observed to form denser fibrin biofilms faster than *S. aureus* 50825. The appearance of initial fibrin strands, assessed visually, was at 1.8 ± 0.3 hr for *S. aureus* 25268 and 2.0 ± 0.0 hr for *S. aureus* 50825. By 24 hr, biofilms reached a mature state for both strains, as indicated by the dense and confluent matrix observed. *S. aureus* 8325-4 developed a fibrin network around the bacterial colonies (Figure S1C), however this formed much later and less dense then the clinical *S. aureus* strains, supporting the fibrinogen utilization data. The tracking fibrin network development showed that fibrin strands did not originate from the bacterial aggregates, but first formed within the plasma appearing to freely move about and then subsequently attach to the aggregates and other fibrin strands (Video 1).

**Figure 6.**
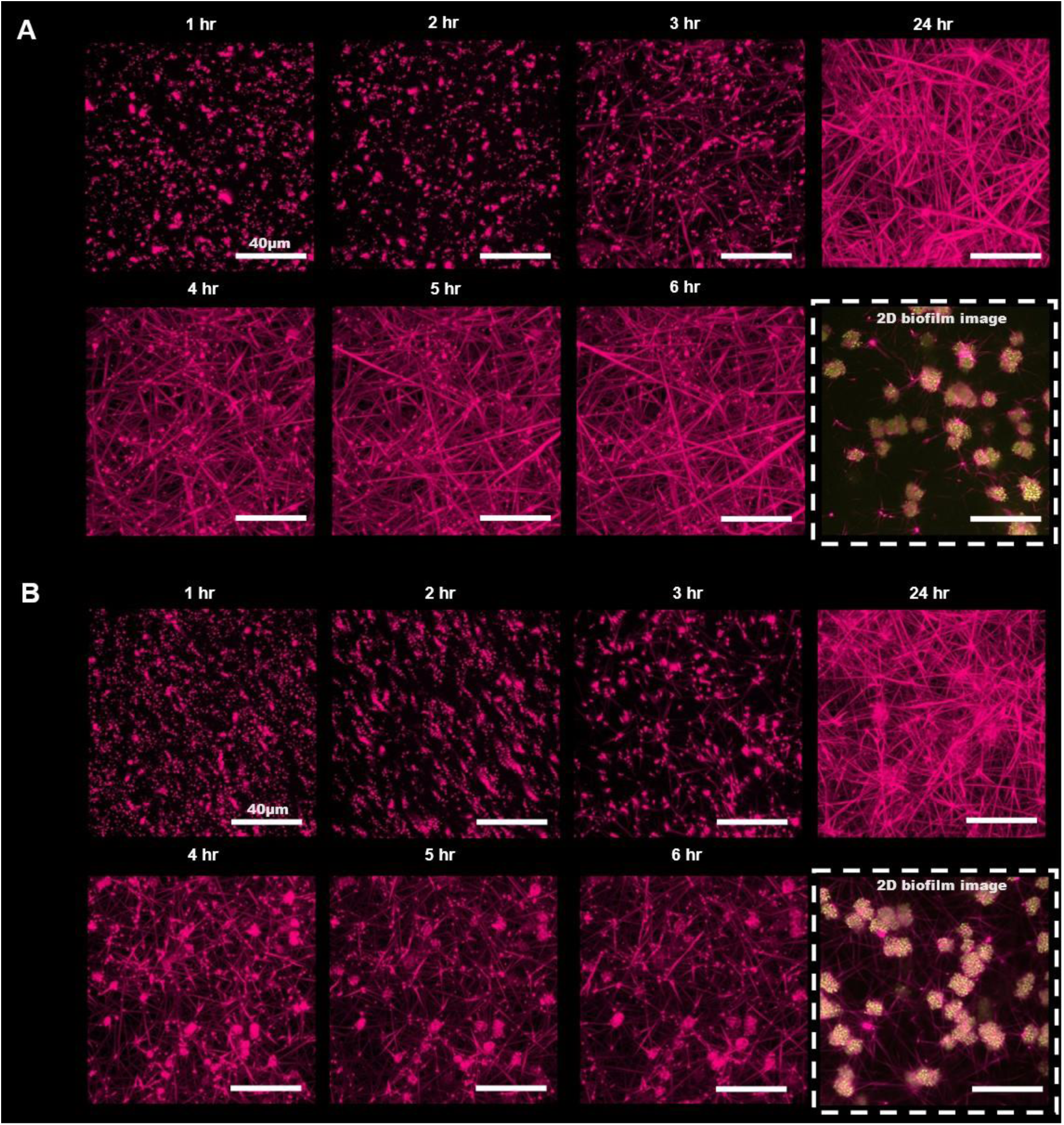
Fibrin network development over time. (**A**, **B**) Three-dimensional volume rendering confocal microscopy of fibrin(ogen) (pink; Alexa Fluor 647) at 1, 2, 3, 4, 5, 6 and 24 hr for two plasma-grown strains, **(A)** *S. aureus* 25268 and **(B)** *S. aureus* 50825. At 24 hr, bacteria were also fluorescently-labeled (yellow; Syto9) to visualize the bacteria within the developed fibrin network, shown as a 2D image (dashed white border). All images are representative of three independent experiments for each strain.

**Video 1.**
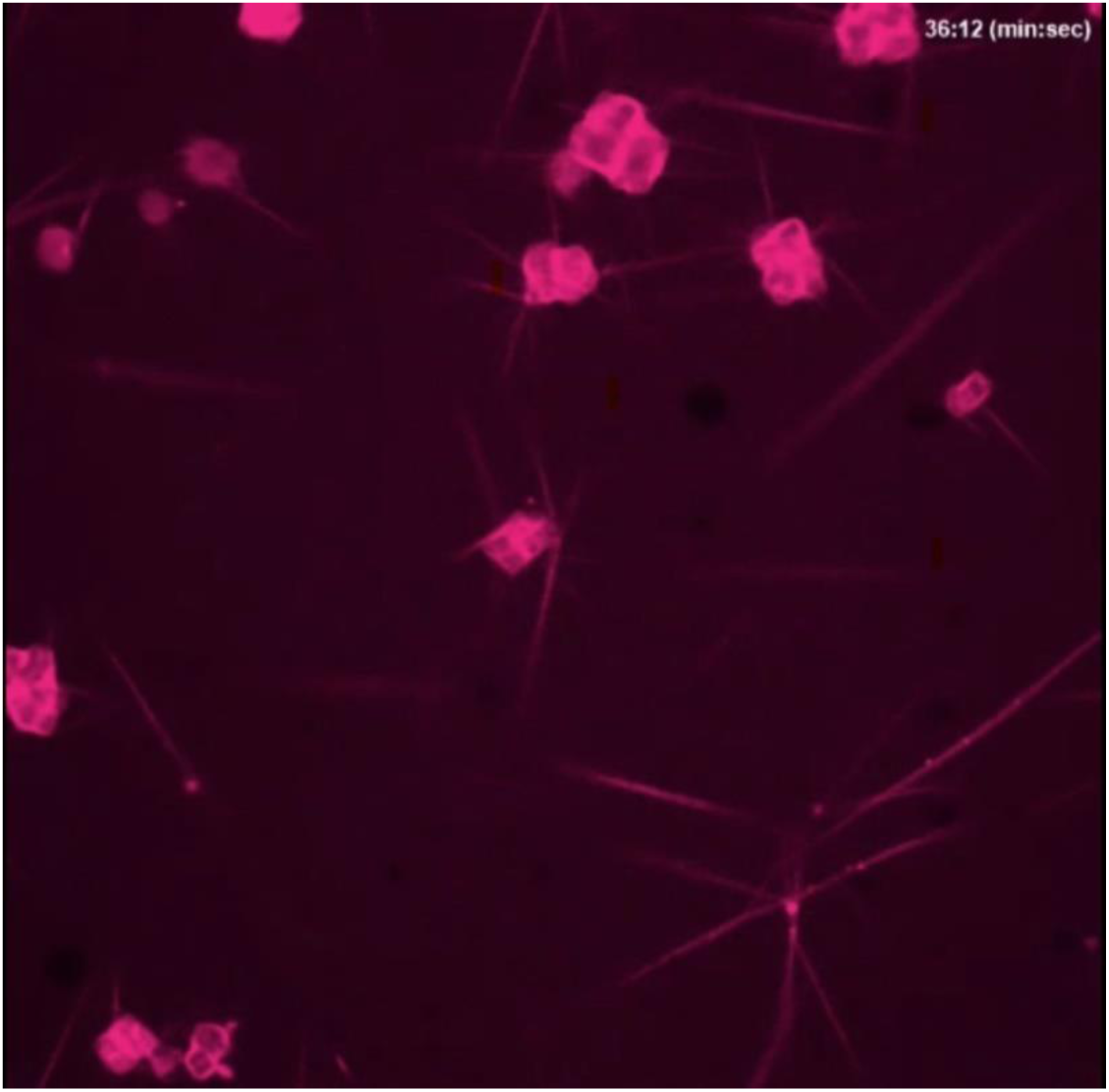
Time-lapse recording of *S. aureus* 25268 fibrin (pink: Alexa Fluor 647) development captured 2 hr and 50 min after inoculation spanning 1 hr and 10 min.

Further analysis of the fibrin network was performed on the images to assess the porosity (Fig. 7A; top graph), and fibrin density (Fig. 7B; bottom graph) of the biofilms over time. Initially, the porosity is high but decreases as the biofilm matures, indicating a denser fibrin biofilm architecture with less free space. *S. aureus* 50825 showed an 82.1 ± 9.0 % decrease (p=0.413) in porosity between 6 and 24 hr, whereas *S. aureus* 25268 showed a 29.7 ± 4.4% decrease (p= 0.003). Despite the variations in porosity between strains, the final porosity of the fibrin network developed by both strains after 24 hr was similar (11.78% difference). The total fibrin(ogen) density in the biofilms (Fig. 7B) increased over time, with a noticeable difference between the two strains. *S. aureus* 25268 produced a more rapid and higher increase in biofilm density compared to *S. aureus* 50825, suggesting strain-specific differences in biofilm formation capabilities. Between 2 and 6 hours, *S. aureus* 25268 showed a 15.0 ± 7.5-fold increase (p= 0.043) in fibrin(ogen) density, and *S. aureus* 50825 showed a 12.2 ± 4.2-fold increase (p= 0.006). This outcome is consistent with the bacterial utilization of fibrinogen, where a significant decrease in fibrinogen levels was found between 2 and 6 hr of biofilm development.

**Figure 7.**
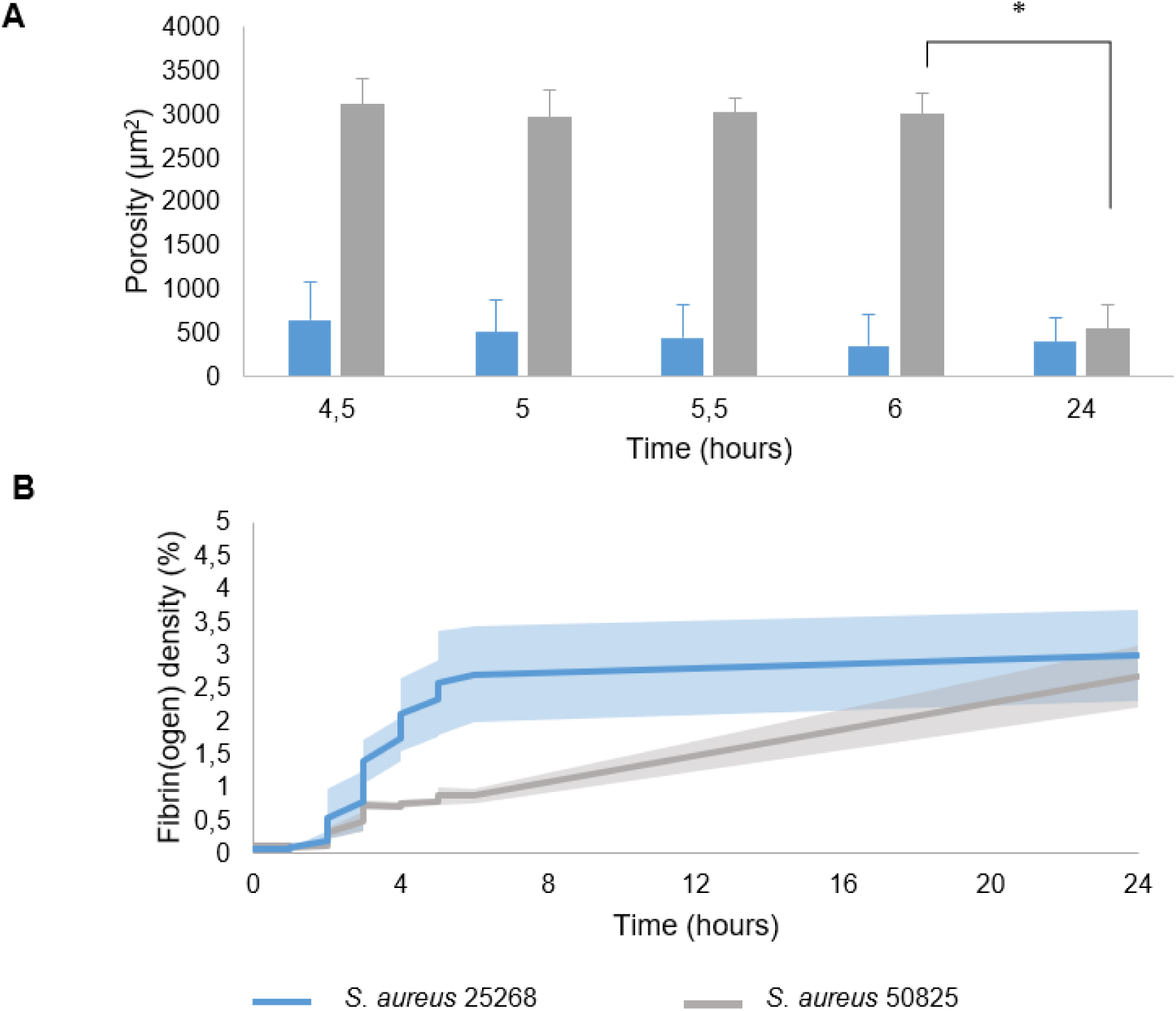
Quantification of the porosity (**A**) and total fibrin(ogen) density (**B**) of the fibrin networks in 3D volume rendering time-lapse confocal microscopy (examples in Fig. 6) of plasma grown *S. aureus* 25268 (blue) and *S. aureus* 50825 (grey). Solid lines represent the mean and shading the standard deviation of three independent experiments.

## Discussion

In this study, we found that bacteria behaved differently depending on the culture medium and substrate, i.e., whether it grew in on porcine valve tissue or hard surface, highlighting the importance experimental design. Only coagulase-producing *S. aureus* isolates formed fibrin biofilms by converting fibrinogen. Furthermore, quantification of the fibrin density and porosity showed strain-dependent fibrin network development over time. These results provide important insights into the significance of coagulase as key mediator of fibrin development in biofilms.

Both bacterial growth and metabolic activity were found to be medium-dependent. For *S. aureus* 25258, maximum metabolic activity was higher in TSB; however, the highest OD was measured in plasma after 24 hr, suggesting a dense biofilm was produced in plasma even though the bacteria were less metabolically active. This difference in maximum metabolic rate can be attributed to the bacterial-specific high nutrient content of TSB, especially its higher glucose levels compared to plasma (2.5 g/l in TSB versus 1.0-1.4 g/l in human plasma [28, 29]. The higher OD in human plasma can be explained by the formation of a dense fibrin biofilm by *S. aureus* (Fig. 4), which is not possible in TSB. Bacterial growth of the coagulase-negative strains was similar across all three media, however their time to reach maximum OD differed with each media. Larger differences were found when comparing to metabolic activity. *S. gordonii* showed limited bacterial growth when assessed using the classical bacterial growth OD assay, while these bacteria were found to be metabolically active. While metabolic activity is a proxy for bacterial growth, it is not synonymous; metabolic activity can also refer to other bacterial activities, such as virulence factor production, indicating that the bacteria are alive and active [30]. Additionally, a slower growing rate of *S. gordonii* was also observed macroscopically, requiring two nights at 37 °C to grow a sufficient number of bacterial colonies on plates for experiments, while *S. aureus* isolates only required one night. Substrate-dependent alterations in metabolic activity were also observed between bacteria growing on heart valve tissue (to mimic infective endocarditis) or a hard surface (to mimic a cardiovascular device infection) (Fig. 2,3), suggesting that the same bacterial strain behaves differently depending on whether it grows on tissue or a device. For example, the tissue surface increased bacterial activity for *S. aureus* 25268 in plasma. Overall, these findings underscore the importance of substrate and medium choice to best match the research study goals, such as the development of soft-tissue infections versus device-related infections and where within or on the human body.

Several *in vitro* and *ex vivo* models have been used to investigate *S. aureus* fibrin biofilm formation [18, 19, 21] such as the classical static biofilm assay [31, 32] and wound infection models [33–35], however, not using 100% human plasma as a growth medium. Models using human plasma provide a more realistic mimic of *in vivo* cardiovascular biofilms compared to traditional growth (e.g., TSB, BHI) or mammalian cell culture media (e.g., RPMI, IMDM), as highlighted by our findings on bacterial responses to different media (Fig.1-3). Coating surfaces and supplementing media with human plasma had been shown to influence biofilm formation and facilitate bacterial attachment [18, 19, 21, 36]. It has also been shown that expression of microbial surface components recognizing adhesive matrix molecules (MSCRAMMs), which are surface proteins bacterial adhesins expressed by *S. aureus* that are essential for initial bacterial attachment, are upregulated in MHB when supplemented with human plasma [19]. We found that coagulase production is crucial for fibrin network formation in *S. aureus* biofilms in human plasma. This finding aligns with prior research highlighting the significance of coagulase enzymes produced by *S. aureus* [21, 37]. Trivedi et al. (2018) studied the role of both coagulases, staphylocoagulase and von Willebrand factor binding protein, by comparing clot generation between coagulase mutant strains [37]. They found that staphylocoagulase mutant strains unable to produce staphylocoagulase could not induce clot formation in a wound like model while a von Willebrand factor binding protein (vWbp) mutant strain could still induce clot formation after 24 hr incubation [37]. Similarly, Zapatozcoa et al. (2015) showed that, on plasma-coated surfaces, a vWbp mutation had no significant effect on biofilm development in RPMI medium [21].

In our study, confocal microscopy imaging of the biofilms after 24 hr revealed fibrin-encased bacterial aggregates in coagulase-positive *S. aureus* strains grown in human plasma (Fig.4). In contrast, coagulase-negative strains showed bacteria on the porcine valves without fibrin. For coagulase-positive *S. aureus* strains, the presence of fibrin provides a physical scaffold for bacterial cells [38], influencing cell-to-cell interactions and promoting the aggregation of cells within the biofilm matrix [39], while other factors, such as complement activation [40], may contribute to the pathogenesis of coagulase-negative strains.

Further investigation into the fibrin network development revealed that the volume of fibrin increases exponentially within the first 6 hr of biofilm development (Fig.6). During this period, fibrinogen utilization by the *S. aureus* 25268 strain increased exponentially (Fig. 5). This finding aligns with a previous study by Rogers (1954), which showed that the coagulase concentration, measured by clotting time in whole human plasma, in culture supernatant increased exponentially within the first 6 hr of development and ceased earlier than bacterial growth [41]. This suggests that coagulases are used primarily during the initial development of the biofilm to form a fibrin biofilm, especially since not all of the fibrinogen in the plasma was utilized (Fig.5). Distinct variations in porosity and density of the developed fibrin networks were observed between *S. aureus* strains (Fig.7). These differences in the fibrin network development could have significant implications for disease progression. One possibility is that the different amount of activity or rate of coagulase produced by *S. aureus* may lead to modifications in the fibrinogen-to-fibrin conversion process, thereby affecting the structural integrity and organization of the resulting fibrin network. These differences may have an impact on disease severity, as Cheng et al. (2010) previously found that time to death in mice was significantly delayed when infections were induced using a coagulase negative mutant *S. aureus* strain instead of a coagulase positive *S. aureus* strain [42].

Human plasma is a better mimic of *in vivo* cardiovascular biofilm development than TSB or IMDM, several factors in the human body, such as immune and red blood cells, are not present in our current model. For instance, fibrin formation can trigger immune activation and subsequent recruitment of inflammatory cells, such as neutrophils, to the infection site, which may influence biofilm development [43]. Moreover, platelets have been shown to contribute to the pathogenesis of *S. aureus* infections by promoting bacterial adhesion and biofilm formation [44]. Therefore, future studies should consider expanding the complexity to further include whole blood components in order to understand how these factors also effect early fibrin biofilm development. Another limitation of this study is the detection limit of 0.4 g/l of fibrinogen in the Sysmex CS-5100 system.

In conclusion, this study highlights the critical role of coagulases in the formation of fibrin biofilms by *S. aureus*. Our findings reveal significant differences in fibrin network development, porosity, and volume between coagulase-positive and coagulase-negative strains, emphasizing the complex interplay between bacterial metabolism and biofilm architecture. The ability of *S. aureus* to utilize fibrinogen and form dense biofilms in human plasma demonstrates the importance of coagulase as a key mediator in the pathogenesis of cardiovascular infections. These insights provide support for novel therapeutic approaches targeting coagulase activity to potentially disrupt biofilm formation and mitigate infection severity [45, 46]. Furthermore, this study showed that the media and even the substrate bacteria grow on influence bacterial activity, emphasizing the need for advanced *in vitro* models that incorporate total human blood components and *in vivo* found substrates. Future research with a focus on understanding the specific mechanisms underlying differences in fibrin network development can provide insights into the pathogenesis of *S. aureus* infections. By advancing our understanding of these critical processes, we can develop more effective strategies to address life-threatening fibrin biofilm-associated infections.

## Author contributions

Conceptualization, S.O. and K.L.; methodology, S.O., G.K., M.M., K.K., and K.L.; validation, S.O., W.W., M.M., K.K., and K.L.; formal analysis, S.O.; investigation, S.O.; resources, W.W., M.M., K.K., and K.L; software, G.C. and J.S.; data curation, S.O., J.H., and K.L.; writing—original draft preparation, S.O.; writing—review and editing, S.O., J.H., M.L.G, G.C., J.S., G.K., W.W., M.M., K.K., and K.L.; visualization, S.O. and K.L.; supervision, K.K. and K.L.; project administration, S.O., and K.L.; funding acquisition, K.K. and K.L. All authors have read and agreed to the published version of the manuscript.

## Acknowledgments

The authors would like to thank Debby van Priem, Iris van Moort, and other members of the Hemostasis Laboratory from the Department of Hematology at Erasmus MCMC, Rotterdam, the Netherlands, for their skillful assistance with the fibrinogen utilization determination. The authors also thank the members of the Therapeutic Ultrasound Contrast Agent group (Biomedical Engineering, Dept. of Cardiology) and the *S. aureus* working group (Dept. of Medical Microbiology and Infectious Diseases) at the Erasmus MC, Rotterdam, the Netherlands for their input, help, and useful discussions.

## Institutional Review Board Statement

Ethical review and approval were waived for this study due to anonymization and de-identification of the bacterial isolates, according to institutional policy.

