## Supplemental Figure 1 for "Early Fibrin Biofilm Development in Cardiovascular Infections"

Supplemental Data:
Manuscript “Early Fibrin Biofilm Development in Cardiovascular Infections”

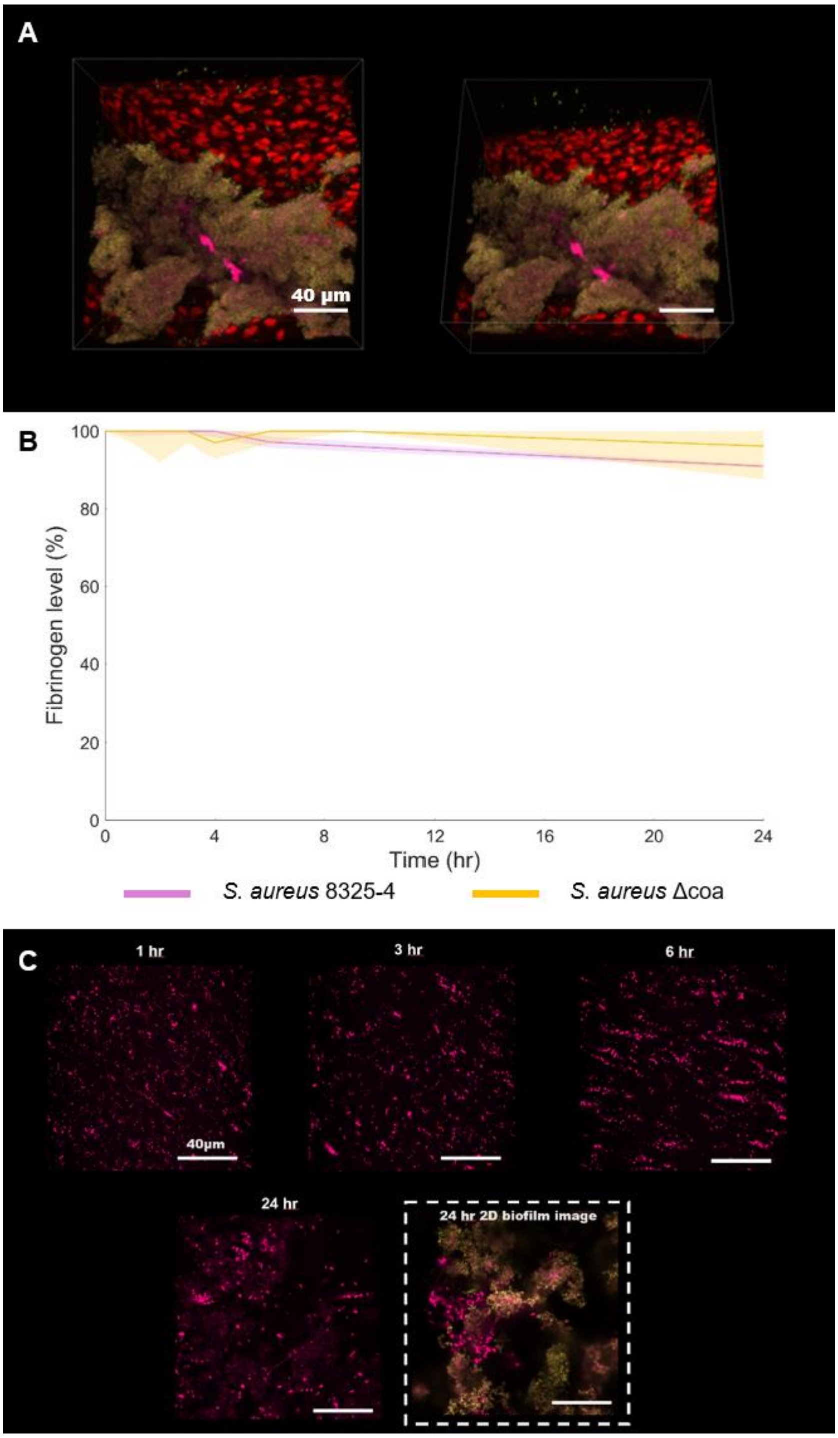


**Supplemental figure S1.** Confocal microscopy and fibrinogen utilization of lab strain *S. aureus* 8325-4. **(A)** 3D volume rendered images (top-view left, angle-view right) of biofilm grown for 24 hr at 37 °C in human plasma on porcine valves, with yellow (Syto9) indicating live bacteria, red (propidium iodide) the valvular cells and dead bacteria, and pink (Alexa Fluor 647) the fibrin and fibrinogen. (**B)** Bacterial fibrinogen utilization of *S. aureus* 8325-4 (purple) and *S. aureus* △coa (yellow) over 24 hr. Solid lines represent the mean and shading the standard deviation. (**C**) 3D volume rendering of fibrin network development over time. At 24 hr, bacteria (yellow) were also fluorescently-labelled (Syto9) to visualize bacteria within the fibrin network. All representative images and data are of three independent experiments.
